# Frequent transitions from night-to-day activity after mass extinctions

**DOI:** 10.1101/2023.10.27.564421

**Authors:** Maxwell E.R. Shafer, Annika L.A. Nichols, Alexander F. Schier, Walter Salzburger

## Abstract

Why certain species survive large-scale extinction events while others do not is poorly understood. While the fossil record can provide insights into morphological adaptations that increase survival probabilities, it provides limited information regarding behavioural traits. Here we integrate behavioural data with phylogenetic comparative models to study the evolution of day-night activity patterns (nocturnality and diurnality) in bony fishes, which have persisted through the last four mass extinction events. Our findings in fish, combined with data across all other clades of vertebrates, provide four lines of evidence that nocturnality conferred an evolutionary advantage during mass extinctions, and that frequent nocturnal-to-diurnal transitions facilitated post-extinction diversification: First, phylogenetic reconstruction indicates that the last common ancestors of all vertebrates and of clades of bony vertebrates were nocturnal. Second, in the lineage of bony fishes, which contains over half of all vertebrate species, twice as many transitions between nocturnal and diurnal activity patterns have occurred compared to tetrapods. Third, within a specific ecological niche, different species exhibit distinct temporal activity patterns, suggesting widespread temporal niche partitioning. Fourth, independent bursts in night-to-day transitions followed large-scale extinction events during the last two geological eras in all four major bony vertebrate groups. These observations suggest that ancestral nocturnality and frequent transitions to diurnality helped vertebrates survive and diversify in the face of extinction events, such as those expected during the current “6th mass extinction” event caused by anthropogenic climate change.

## INTRODUCTION

The Earth’s geological timeline has repeatedly been punctuated by mass extinctions^2^. Such events are typically associated with climate change^3^ and macro-habitat turnovers^4^, leading to the downfall of many species and a substantial decline in organismal diversity. On the other hand, surviving lineages may undergo extensive diversification subsequent to extinction events, often facilitated by ecological opportunity in the form of vacated niches previously occupied by extinct species^5^. In animals, it is becoming increasingly clear from the fossil record that survival probabilities during mass extinctions are influenced by biotic factors such as body size^6^ and physiology^7^, as well as by distribution range^8,9^, albeit to varying degrees depending on the clade^10^. However, other potentially important determinants of survival probability, including behaviours and life history traits, are difficult to determine from fossils alone. Ancestral state reconstructions and phylogenetic modelling can offer insights into these traits. For example, applying such approaches has revealed that non-arboreal lifestyles were selectively advantageous in birds^11^ and mammals^12^ during the rapid deforestation across the globe associated with the Cretaceous-Paleogene (K-Pg) mass extinction event^11^.

Animal species exhibit characteristic diurnal or nocturnal activity patterns. It is well established that these distinctive circadian rhythms are the result of adaptations to the daily light cycle^13^. Much less is known about the contribution of diurnal versus nocturnal behaviours to macro-evolutionary patterns such as diversification and extinction. For example, what is the role of nocturnality or diurnality during selection caused by mass extinction events? And how do evolutionary shifts between nocturnality or diurnality contribute to diversification following these events? Previous studies have suggested that early mammals’ nocturnality minimised competitive overlap with diurnal dinosaurs^14^. After the K-Pg boundary and extinction of non-avian dinosaurs, mammals diversified into diurnal niches^15^. These and other studies in tetrapods^19^ suggest that temporal niche partitioning, wherein species avoid competition by occupying different temporal niches^13^, might be involved in survival during, or diversification after, mass extinction events. However, it remains unclear how vertebrate diurnality and nocturnality evolved over long time periods and how diurnal-nocturnal transitions affected species survival and diversification.

Here we report the first large-scale examination of the evolution of temporal activity patterns across all vertebrate groups with a particular focus on bony fishes, which constitute more than half of all vertebrate species and have persisted through four mass extinction events. Analysis of extensive datasets of day-night activity patterns^15,19^ and hidden Markov Models of trait evolution indicate that the last common ancestor of all vertebrates and the ancestors of all bony vertebrate groups (bony fish, amphibians, mammals, and sauropsids) were nocturnal. Notably, in fishes, we observed twice as many transitions between diurnality and nocturnality compared to tetrapods when accounting for overall diversity across lineages. We link bursts in nocturnal-to-diurnal transitions to geological boundaries and associated climate shifts and increased extinction rates. Our results suggest that ancestral nocturnality conferred an evolutionary advantage during mass extinctions, and that the post-extinction diversification of fishes was facilitated by the frequent day-night activity transitions that led to temporal niche partitioning (nocturnal versus diurnal). We speculate that nocturnality might have a selective advantage during current and future large-scale extinction events.

## RESULTS

### A nocturnal ancestor for bony and cartilaginous fishes

To examine the evolution of temporal activity patterns across fishes we first compiled a dataset comprising information on 3,985 species of bony and 135 cartilaginous fish (**Supplementary Data**), covering 96% of all orders, 68% of families, 31% of genera, and 12% of known fish species (https://www.fishbase.se/, version (06.2021)) (**Extended Data Figure 1a-b**). Based on the available literature, we assigned each species into four temporal activity patterns categories: diurnal (greater activity during the day), nocturnal (greater activity during the night), crepuscular (peaks of activity during the dawn or dusk), or cathemeral (irregular/arrhythmic activity patterns, or no difference between night/day/dawn/dusk) (**Extended Data Figure 1c**). In our database, species can be both crepuscular as well as diurnal or nocturnal if they had peaks of activity during dawn/dusk, but also higher activity during the day or night periods. The majority of entries (∼70%) come from primary literature sources directly measuring fish abundance, metabolism, or activity in the field or in the lab (**Extended Data Figure 1**). However, many studies do not contain precise enough temporal resolution to quantify crepuscular and cathemeral activity patterns. Therefore, for the majority of our analyses, we focused on only those species that were assigned as either diurnal or nocturnal.

We next reconstructed the likely temporal activity pattern at ancestral nodes across a composite phylogenetic hypothesis of all diurnal and nocturnal fish species^16^ (**Figure 1a** and **Extended Data Figure 2-3**). These methods have been used to reconstruct the evolutionary history of behaviours^11,12^, including diurnality and nocturnality across mammals and tetrapods^15,19^. Comparisons between different models favoured scenarios with hidden transition rates over simpler models with single transition rates^17^ (**Extended Data Figure 2a**). Hidden rate models allow for heterogeneity in the transition model, and for transition rates to vary with time. Specifically, using hidden rates allowed for two versions of each state (diurnal and nocturnal), each with different transition probabilities between states.

**Figure 1.**
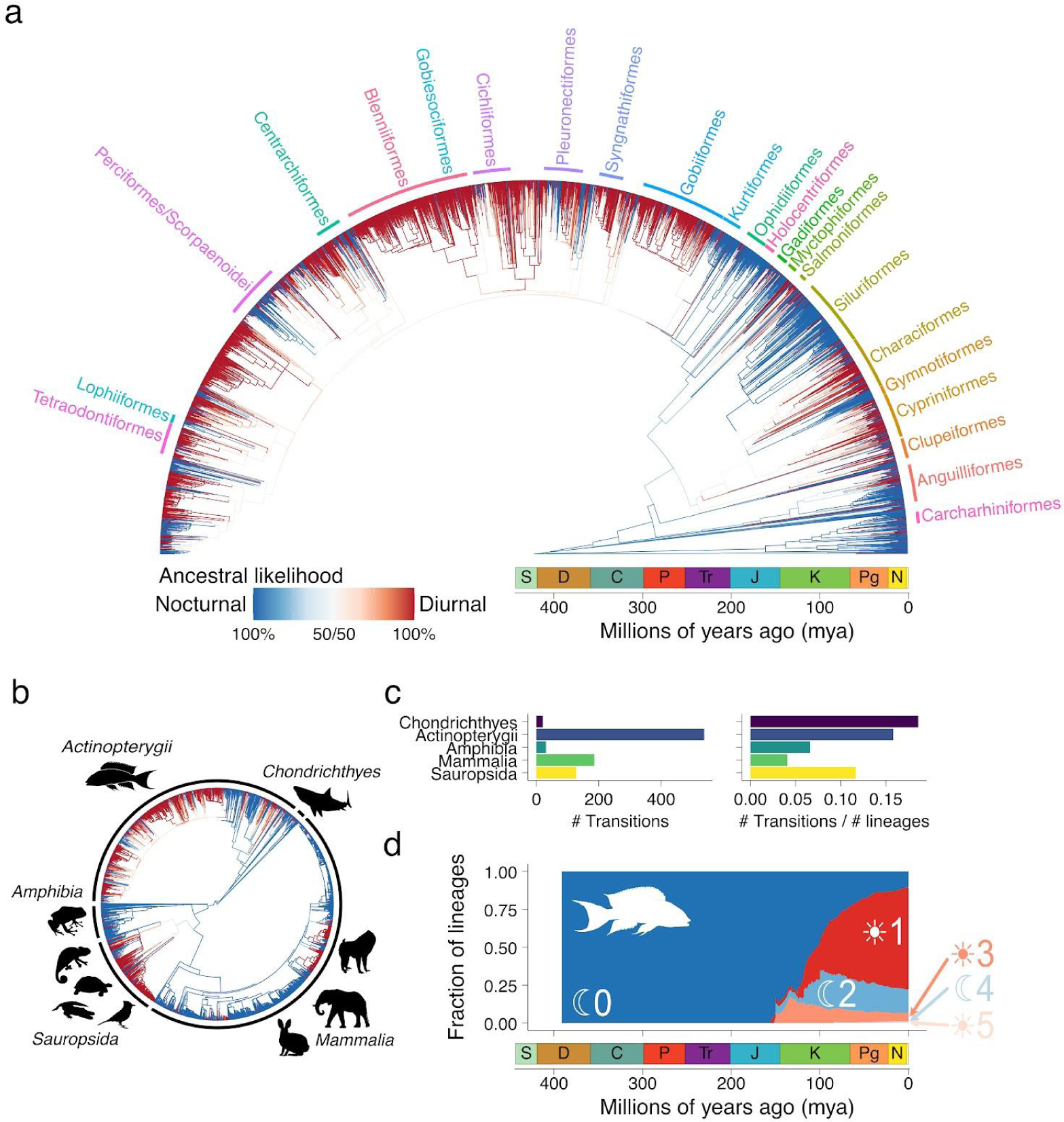
Evolution of diurnal/nocturnal behaviour across vertebrates. **a**) Marginal ancestral reconstruction across bony and cartilaginous fishes with major Orders labelled (3524 species). **b**) Marginal ancestral reconstruction across all vertebrates. Branches and tips are coloured by their likelihood of being diurnal (red) or nocturnal (blue). **c**) The total number of transitions between diurnal and nocturnal behaviour, and the number of transitions normalised by the total number of lineages in each dataset, across five major vertebrate groups. **d**) The accumulation of lineages across time in bony fish with a reconstructed history of 0, 1, 2, 3, 4, or 5 transitions between diurnal/nocturnal behaviour. Data on tetrapods comes from Anderson & Wiens 2017, and data on mammals from Cox et. al., 2021.

We found that the best fitting scenario for the evolution of temporal activity patterns in fishes strongly supports nocturnality in the last common ancestors (LCAs) of both bony and cartilaginous fish, and at least two transition rate classes (**Figure 1a** and **Extended Data Figure 2b**). In both fish groups, temporal activity pattern evolution is characterised by extensive transitions between diurnality and nocturnality, with one hidden rate class having high transition probability of both diurnal to nocturnal and nocturnal to diurnal switches (fast rate), and a second hidden rate class having transition probabilities of essentially zero (slow rate) (**Extended Data Figure 2b**). Indeed, models where we enforced one rate class to be incapable of transitioning had only marginally worse likelihood than ones where rates were unconstrained (**Extended Data Figure 2a**).

These results suggest that lineages are either in a state that is incapable of transitioning between temporal activity patterns or are in a state that is capable of transitioning between states and do so frequently. As examples of lineages likely incapable of transitioning, virtually all butterflyfish (*Chaetodontidae*) and blennies (*Blenniformes*) are diurnal, whereas almost all catfishes (*Siluriformes*), squirrelfish (*Holocentriformes*), and cardinal- and nurseryfish (*Kurtiformes*) are nocturnal. Many other clades were composed of high numbers of both diurnal and nocturnal species, including the hyperdiverse cichlids (*Cichliformes*), bottom-dwelling flatfish (*Pleuronectiformes*), and the cryptic scorpion- and rockfish (*Scorpaenoidei*) (**Figure 1a**). Eels (*Aquilliformes*) contain nocturnal and diurnal species, but also a large number of arrhythmic species (**Figure 1a**). Interestingly, high transition probabilities between temporal activity patterns (fast rate) were primarily observed in lineages with high diversification rates, including the *Characiformes* and *Cypriniformes*, as well as the *Acanthomorpha*, a clade that contains approximately one quarter of all vertebrate species^18^ (**Figure 1a**). These patterns were consistent whether or not our analyses included both bony and cartilaginous fishes, either group alone, or only species whose activity patterns were measured in the field or the lab (**Extended Data Figure 3b-d**). Overall, these results suggest that numerous clades within fishes independently evolved the ability to frequently transition between temporal activity patterns, whereas others have conserved their temporal activity patterns for millions of years.

### Transitions between activity patterns are remarkably dynamic in fishes

Fishes are not a monophyletic group, as bony fish are the sister group to tetrapods, which include the amphibians, mammals, and sauropsids (reptiles, birds). Bony fish and tetrapods together are the sister group to cartilaginous fishes. A recent landmark study has shown that the last common ancestor of tetrapods was nocturnal^19^. However, unlike the fish groups in our study, temporal activity patterns within tetrapods are evolutionarily conserved, with only a few transitions away from ancestral nocturnality^19^. Combining our database of fish temporal activity patterns with data from two recent studies covering mammals^20^ and all tetrapods^19^, and performing ancestral state reconstructions on this larger dataset, revealed that the last common ancestor of all vertebrates was also most likely nocturnal (**Figure 1b**).

To quantify the minimum number of transitions between temporal activity patterns that have occurred in different clades, we used the probability of each state at each internal node of the phylogeny from our marginal ancestral state reconstructions. We assigned a temporal activity pattern (diurnal or nocturnal) to all internal nodes of each tree based on the most probable state. We then determined for each node if it represented a transition in temporal activity pattern from its parental node. This analysis was performed on each of the four major groups of bony vertebrates (bony fish, amphibians, mammals, and sauropsids) using independent ancestral state reconstructions for each group (**Extended Data Figure 3-4**). In all clades, models with two hidden rates outperformed single-rate models, and were characterised by one rate class with a high transition probability and one with a very low transition probability (**Extended Data Figure 3-4**). In the case of sauropsids, the fast rates were not paired, that is, rate class 1 was characterised by a high probability of transitioning from diurnal to nocturnal and a low probability from nocturnal to diurnal (rate class 2 was the opposite).

We identified 538 transitions across bony fish and 21 within cartilaginous fish, representing 0.15 and 0.18 transitions per extant lineage in our trees, respectively (**Figure 1c**). Tetrapods, on the other hand, had less than half the number of normalised transitions compared to fish (345 transitions, 0.06 per lineage), but patterns in transitions were disparate between clades. The lowest rates were observed in mammals (186 transitions, 0.04 per lineage) and amphibians (31 transitions, 0.07 per lineage), with much higher rates in sauropsids (128 transitions, 0.12 per lineage). However, transitions in sauropsids occurred almost entirely in the direction from nocturnal to diurnal (72% of extant lineages characterised by a single transition), with rare reversals to nocturnal behaviour in birds and legless lizards (genus *Calyptommatus)* (**Extended Data Figure 3c**). In contrast, the evolutionary lineages leading to 90% of extant fish species have undergone at least one transition, 28% transitioned at least twice, 8% three times, and 2% four times (**Figure 1c-d**). There were even 0.5% of extant fish species that may have an evolutionary history characterised by five or more transitions between nocturnality and diurnality over the past 430 my (**Figure 2d** and **Extended Data Figure 5**). Species of toothless characins (*Characiformes*) and rockfish (*Scorpaeniformes*) including *Sebastes serranoides* and *S. mentella* were among those with a history of more than four transitions. Therefore, whereas tetrapods are characterised by evolutionary conservation in temporal activity patterns, bony fish are characterised by evolutionary lability.

**Figure 2.**
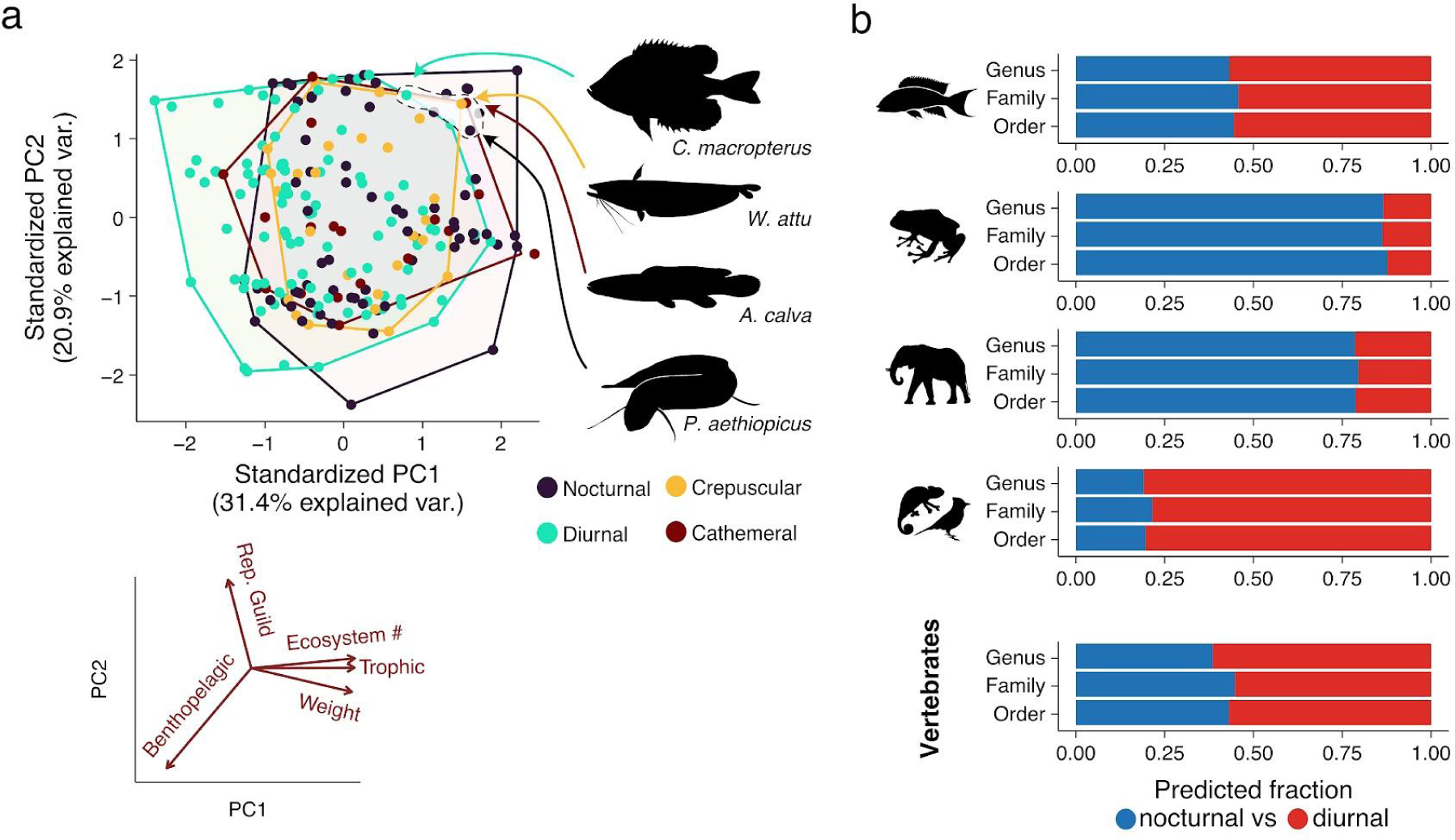
Overlap in ecological niche space across temporal niches in fish. **a**) Principal component analysis across five ecological traits in bony fishes and the associated loadings across traits (reproductive guild, ecosystem number, trophic level, mean weight, benthopelagic axis). Example species and their location in 2-dimensional PCA space of nocturnal, diurnal, crepuscular, and arrhythmic species with highly similar ecologies. **b**) Predicted numbers of total diurnal and nocturnal species for different taxonomic levels for bony fish, amphibians, sauropsids, mammals, and all vertebrate species.

### Ecological niche spaces are consistent with temporal niche partitioning

The high numbers of both diurnal and nocturnal species of fish raised the possibility of extensive temporal niche partitioning: species in the same ecological niche might avoid competition by changing when they are active^13^. Alternatively, expansion of clades into different temporal niches could be limited by ecological factors. For example, the ecologies of nocturnal and diurnal mammalian species are only partially overlapping, suggesting that different abiotic and biotic selective forces are in play during the day versus during the night^20^.

To test whether or not temporal niches are constrained by species’ ecology, we developed an ecological economics spectrum for fish^21^. This spectrum compiles data on a fish species’ body mass, investment in reproduction, trophic level, bentho-pelagic habitat, and ecosystem diversity, and can be reduced to a two-dimensional hypervolume through principal component analysis (PCA), similar to what has been done previously in mammals^20^. Comparison of ecological hypervolumes between diurnal, nocturnal, crepuscular, and cathemeral fish species revealed a strong overlap between groups, with no hypervolume space being specific to any temporal activity pattern (**Figure 2a**). Distributions across individual ecological traits also showed no differences between temporal activity patterns (**Extended Data Figure 6**). For example, the nocturnal *Protopterus aethiopicus*, the diurnal *Centrarchus macropterus*, the crepuscular *Wallago attu*, and the arhythmic *Amia calva* are all medium to large invertivores/piscivores that live in multiple habitats (rivers, lakes, lagoons, sea bays) and build and guard their egg nests (**Figure 2a**). These results suggest that in contrast to mammals, for every ecological niche, there are fish species occupying different temporal versions (diurnal and nocturnal) of that same ecological niche.

Extensive temporal niche partitioning also predicts that globally there should be similar numbers of diurnal and nocturnal species of fish. To test this hypothesis we used the proportions of diurnal, nocturnal, crepuscular, and cathemeral species across taxonomic levels in our database to predict global proportions across lineages based on total numbers of species in each taxonomic level. Our database contains 2,084 diurnal (54%), 1,183 nocturnal (30%), 308 crepuscular (8%), and 314 cathemeral (8%) fish species (**Extended Data Figure 1** and **Extended Data Figure 7**). However, our extrapolations and predictions for the total number of diurnal or nocturnal fish species were remarkably similar (55% diurnal and 45% nocturnal) (**Figure 2b** and **Extended Data Figure 7**). This result agrees with studies in our database relying on high resolution tracking information (categories D and E), which report an even closer distribution between diurnal (47%) and nocturnal (53%) fish species. This analysis suggests that the literature is overrepresented by references to diurnal clades, such as the well-studied diurnal reef fishes, while missing high resolution information from nocturnal orders like ghost sharks (*Chimaeriformes*), swamp eels (*Synbranchiformes*), electric knifefish (*Gymnotiformes*), and electric rays (*Torpediniformes*) (**Extended Data Figure 7c**). Combining our predictions in fish with those for all other tetrapod groups resulted in an relatively even distribution between diurnal and nocturnal species across all vertebrates (57% and 43%, respectively) (**Figure 2b**), demonstrating that temporal niche partitioning is a vertebrate-wide phenomenon: for every ecological niche there is likely both a diurnal and a nocturnal species, and that missing counterparts of nocturnal mammals would be likely found within the diurnal sauropsids.

### Transitions in activity patterns are associated with mass extinction events

Our analysis of the evolutionary history of activity patterns across all four major bony vertebrate groups reveals large scale temporal niche partitioning that is best explained by models with two transition rates: one which allowed transitions between states (fast rate) and one that did not (slow rate) (**Extended Data Figure 2a** and **Extended Data Figure 3-4**). Do the fast transition rates correspond to specific paleo-environmental conditions or events that triggered temporal niche transitions, and thus allowed those clades to partition temporally? To test this idea, we compared the rates of transitions between groups and across 430 my of evolution.

The expansion from ancestral nocturnality and temporal activity pattern evolution in bony vertebrate groups displayed similar dynamics over evolutionary timescales (**Figure 3a**). In both sauropsids and bony fish we observed a burst of transitions from nocturnality to diurnality beginning around 140 mya, followed by variable and decreasing transition rates over the last 140 my (**Figure 3a**). Our analysis confirms the more recent burst of transitions from nocturnal to diurnal activity in mammals approximately 65 mya^15^, and demonstrates a similar peak in amphibians (**Figure 3a**). We also observed many transitions from diurnality to nocturnality in bony fish over the last 130 my, and some in the other tetrapod lineages over the last 30 my, though these were fewer in number compared to nocturnal to diurnal transitions (**Figure 1d**, **Extended Data Figure 4** and **Extended Data Figure 8a**). For all four vertebrate lineages (bony fish, amphibians, sauropsids, and mammals) our models do not predict transitions or diurnal lineages before these bursts and before 140 mya.

**Figure 3.**
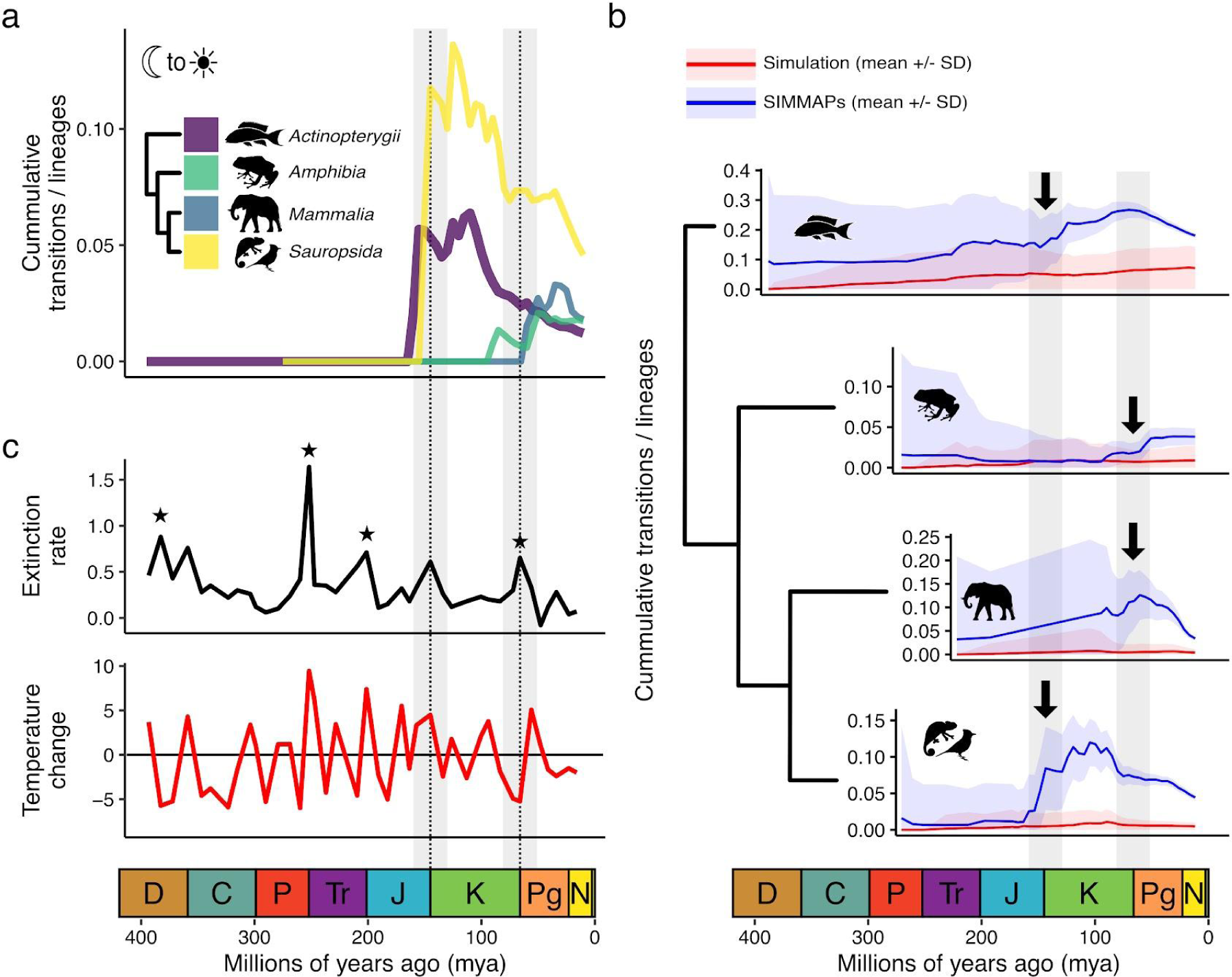
Nocturnal bottlenecks across bony vertebrates are associated with paleontological boundary periods. **a**) Cumulative number of transitions from nocturnal to diurnal activity patterns divided by the cumulative number of lineages per bony vertebrate group. Inset: the phylogenetic relationship between the groups. **b**) Comparisons of 500 simulations of the evolution of diurnal/nocturnal behaviour with 500 samples from the posterior distribution of our ancestral reconstructions (SIMMAPs) across bony vertebrate groups. Arrows indicate when simulations and SIMMAPs begin to differ. Dark lines are the means of simulations or SIMMAPs and shaded areas represent one standard deviation. **c**) Patterns in extinction rate (black line) and temperature change (red line) across the paleontological record. Stars represent ‘mass extinction’ events. Transitions, extinction rates, and temperature changes are averaged across 5 million year bins.

A lack of diurnal states and of transitions before 140 mya suggests two possibilities, 1) that these lineages were nocturnal and incapable of transitioning before these periods, or 2) they were able to transition to diurnality but diurnal species did not survive. To differentiate between these scenarios, we performed simulations of the evolution of temporal activity patterns and compared these simulations to the ancestral reconstructions and transitions derived from stochastic character maps (SIMMAPs). SIMMAPs represent possible histories sampled from the probabilities in our marginal ancestral reconstruction^22^. Simulations are possible histories of temporal activity pattern evolution from the root of the tree, given only the transition rates from our hidden rates models, and not considering tip states. Simulations of temporal activity pattern transitions suggest that there should have been the same or a higher number of transitions in temporal activity patterns (diurnal to nocturnal, and nocturnal to diurnal), in all four groups between 400-140 mya as compared to 140-0 mya (**Figure 3b** and **Extended Data Figure 8**). In contrast to the simulations, stochastic maps display distinct increases in transition rates around 140 mya at the Jurassic-Cretaceous (J-K) boundary for fish and sauropsids, and around 65 mya at the Cretaceous-Paleogene (K-Pg) boundary for amphibians and mammals (**Figure 3b**). This suggests that all four bony vertebrate groups were able to transition before these geological boundary periods, but that only nocturnal lineages survived the respective events.

Boundaries between geological periods are characterised by global changes in temperature and CO_2_ levels, and can be associated with changes in sea level and continental fragmentation^23^. These events can be accompanied by elevated extinction rates, typically in the form of mass-extinction events^3^. The pattern of nocturnal to diurnal transitions in mammals is known to be associated with the mass extinction event that eradicated non-avian dinosaurs at the K-Pg boundary^15^, which could also explain the elevated transition rate we observe in amphibians (**Figure 3a**). Peaks in transition rates within bony fish and sauropsids also had a high correspondence to the elevated extinction rates and biotic upheaval occurring during the J-K boundary^24^ (**Figure 3a**). Patterns of origination from fossil records demonstrate that all four major bony vertebrate lineages existed and diversified extensively before these events, and before extant lineages transitioned away from ancestral nocturnality (**Extended Data Figure 9**). Together, these results suggest elevated extinction rates and major biotic upheaval as triggers for temporal niche transitions, rather than periods of origination or diversification. This analysis suggests that nocturnal lineages were more robust to extinction, potentially outliving diurnal competitors and contributing to the recovery of species diversity during at least two recent geological periods (Cretaceous and Paleogene) through temporal niche partitioning.

In summary, our ancestral reconstruction of temporal niche across vertebrates reveals that the last common ancestor of all vertebrates was nocturnal, and that nocturnal lineages may be more resistant than diurnal ones to the biological upheaval and extinction associated with geological boundary periods. These events were associated with elevated temperatures^27,28^. The current anthropogenic climate change is also responsible for large shifts in temperatures outside of historical norms^29^. Activity during the cooler nocturnal periods may be beneficial during these epochs, providing a mechanism for species survival^30^. Our models predict nocturnality as an advantage for survival under increased temperatures and thus inform current biodiversity efforts to counteract the effects of climate change.

## METHODS

### Data collection/annotation

Tracking of locomotory activity patterns is routine in lab animals and observations of animals in the field across the diel cycle are common. To identify temporal activity patterns for fish species we performed a systematic literature search inspired by the work of Karhl et. al., 2021^31^. We used RISmed v2.3.0 package in R to search pubmed for all possible taxonomic levels (order, family, genus, and species names) in combination with “diurnal OR nocturnal OR crepuscular OR cathemeral OR diel OR circadian” (https://cran.r-project.org/). These were supplemented with searches in Google Scholar for diel activity terms in combination with taxonomies that were missed in Pubmed searches, including all family and genus names from orders that were not already represented in the database. Taxonomies of fish species were derived from Fishbase (https://www.fishbase.se/, version (06.2021)). Data for tetrapods was from Anderson & Wiens 2017^19^, and data from mammals was from Cox et. al., 2021^20^.

Data were categorised (A, B, C, D, or E) based on the type of experiments and analysis done, rather than the quality or quantity of the data. Data sources included entries in public databases without primary literature such as Wikipedia (category A), quantification of diel variation in catch rate (category B), quantification of diel variation in feeding rates (field or lab), or quantification of metabolic rates including diel variation in oxygen consumption or the production of metabolic byproducts (category C), direct observation of animals in situ (transects, visual surveys, video recording), or direct tracking of individuals over diel periods in the field (satellite or sonar tags) (category D) or in a laboratory setting (video recording, observations) (category E).

### Tree construction

To generate a single tree based on a composite phylogenetic hypothesis we used the list of taxa from our database to generate a tree from the Open Tree of Life (v. 14.7, http://opentreeoflife.org) using the R package rtol v3.0.12 and the function tnrs_match_names^16,32^. Since the literature sources for our activity patterns date from over 100 years ago, and because many taxonomic revisions have occurred within fish taxonomic ranks, we allowed approximate matching. Each approximate match was examined for accuracy and we used databases such as Fishbase (https://www.fishbase.se/, version (06.2021)) and the World Register of Marine Species (https://www.marinespecies.org) to compare and correct for taxonomic revisions where possible.

We downloaded time calibrated datasets for our taxa from TimeTree v5^33^. We then used congruify.phylo function from geiger v2.0.10 to time-calibrate our composite phylogenetic tree to add branch lengths^34^. For independent analysis of mammals we use the maximum likelihood tree based on the 10k trees sampled from the posterior distribution available in Cox et. al., 2021^20^. For independent analysis of tetrapod lineages we used the tree from Anderson & Wiens 2017^19^, and subsetted it to include only amphibia or sauropsida. For analyses of all vertebrate species, we generated a synthetic phylogenetic tree using the Open Tree of Life using the taxa from the combined datasets (mammalia, sauropsida, amphibia, and bony and cartilaginous fishes), and time-calibrated using data from TimeTree v5 as we did above for fish.

### Predictions of total temporal niche diversity across clades

To account for potential biases in the literature and our literature survey, and to estimate the overall distribution of temporal niches in fish, we multiplied the fractions of each niche within each clade by the overall species richness for that clade. We did these calculations at the order, family, and genus taxonomic levels, and determined average values across these comparisons. By this estimation, there are approximately 14,700 diurnal (43%), 12,600 nocturnal (37%), 3,400 crepuscular (10%), and 3,100 cathemeral (9%) fish species in the world. To determine predicted numbers for tetrapod groups, we performed the same analysis across taxonomic levels in mammals, amphibians, and sauropsids (reptiles and aves). Mammalian data was derived from the American Society of Mammalogists Mammal Diversity Database (https://www.mammaldiversity.org/, accessed August 17^th^, 2023)^35,36^, amphibian data from AmphibiaWeb (https://amphibiaweb.org, accessed August 17^th^, 2023), reptile data from The Reptile Database (http://www.reptile-database.org, accessed August 17^th^, 2023)^37^, and data on aves from The eBird/Clements checklist of Birds of the World (https://www.birds.cornell.edu/clementschecklist/download/, accessed August 17^th^, 2023).

### Models of character evolution

We tested the fit of Equal Rates (ER), All Rates Different (ARD), and Hidden Rates (HR) models (with 2 or 3 hidden rates) for the evolution of nocturnal vs diurnal character states using the function corHMM from the package corHMM v2.7^17^. The models were compared using both the maximum likelihood and Akaike Information Criterion (AIC). We found that HR models fit the data better than either ER or ARD models, and that the fit increased with the number of hidden rate classes. In general, the interpretability of HR models with more than 2 rate classes decreased, but did not differ substantially from those with 2 rate classes across clades. In all cases, one of the hidden rate classes was characterised by a transition rate of nearly 0, and one or more hidden rate states with high transition rates. Indeed, manually assigning one rate class a transition probability of exactly 0 had no effect on the likelihood or AIC scores compared to an unrestrained model with the same number of rate classes.

### Ancestral state reconstruction

Ancestral character states were reconstructed using corHMM from the package corHMM v2.7. We used marginal ancestral reconstruction to determine the likelihood of each state at each internal node and at the tips for HR models. For HR models, to determine the likelihood of each character state or rate class we summed the likelihoods across multiple states (e.g. the sum of “Diurnal rate class 1” and “Diurnal rate class 2” for “Diurnal” or the sum of “Diurnal rate class 1” and “Nocturnal rate class 1” for “Rate class 1”).

### Simulations and stochastic character maps

Simulations of diurnal/nocturnal character evolution were performed using the rTraitDisc function from ape v5.6-2^38^ using the rates derived from the best fitting model, and the most likely root state from our ancestral reconstructions. To generate SIMMAPs from HR models from corHMM we used the function makeSimmap from corHMM^39^ using the rates from our best fitting model. This is similar to the function make.simmap from phytools^22^. We generated 500 simulations and 500 SIMMAPs for each clade.

### Analysis of transitions through time

To determine the evolutionary history of transitions within clades we extracted the likelihood from the marginal reconstruction for the most recent common ancestors, or all ancestors for each node (using the Ancestors function from phangorn v2.7.0)^40^. Node ages were extracted using nodeHeights from phytools v1.0-3 and used to assign timepoints for each transition^22^. The likelihood from the marginal reconstruction was rounded to the most likely state - e.g. any node with greater than 50% likelihood of diurnal was assigned diurnal. We determined cumulative sums for both lineages and transitions and calculated the proportion of lineages transitioning through time by dividing the cumulative transitions by cumulative lineages at each time point. These numbers were then averaged across 5 my long time bins. This analysis was performed using custom code written in R and is available online (https://github.com/maxshafer/fish_sleep).

The same method was repeated for simulated trait data by identifying parent/daughter node states and transitions for each simulation. Importantly, SIMMAPs allow for transitions to occur along branches (in between nodes) which resulted in more overall transitions compared to analysis using the likelihoods from our marginal ancestral reconstruction (using only node states). This is likely due to the high transition rates, and long branch lengths in our fish tree. However, we identified states at internal nodes for each SIMMAP, and used the same method as for ancestral reconstructions and simulated data to determine lineage transitions through time. This approach allowed for more direct comparison between ancestral reconstructions, simulations, and SIMMAPs, with transition times limited to nodes and node ages in the dataset. In either case, dynamics of transitions through time were nearly identical between analyses, suggesting that the discrepancy is only in the number of transitions, not when they occur. We computed mean values and standard deviations for the transition rates (cumulative transitions divided by cumulative lineages) across 500 simulations and 500 SIMMAPs.

### Collection and analysis of paleological and ecological data

Paleontological data on extinction rates and temperature changes were taken from Song et. al., 2021^3^. Data on fossil records were extracted from the Paleobiology Database (https://paleobiodb.org/). The final data used were downloaded from the Paleobiology Database on 8 August, 2023, using filters for taxonomic class (“Amphibia”, “Mammalia”, “Actinopterygii”, or “Sauropsida”).

Data on the ecologies and habitats of each taxon were acquired from Fishbase (https://www.fishbase.se/, version (06.2021)) using the package rfishbase v4.0^41^. We used information on each species’ weight, trophic level, ecosystem breadth, benthopelagic axis, and reproductive strategy. For weight we used log transformed mean measurements of maximum weights (391 species). Trophic level was derived as a mean of ‘DietTroph’ and ‘FoodTroph’ and ranged from 1-5 (low to high trophic level, 653 species). Ecosystem breadth was calculated as the number of unique ecosystem types (‘lake’, ‘lagoon’, ‘river (basin)’, ‘sea/bay/gulf’) occupied by each taxon and ranged from 1-4 (low to high breadth, 924 species). The benthopelagic axis was derived by converting habitats to a numeric scale (‘demersal’, ‘reef-associated’, ‘benthopelagic’, ‘pelagic-neritic’, ‘pelagic’, ‘pelagic-oceanic’) with ‘demersal’ equalling 1, and ‘pelagic-oceanic’ equalling 6 (923 species). Finally, reproductive strategy was converted similarly, with species ranging across ‘open water/substratum’, ‘egg scatterers’, ‘nonguarders’, ‘brood hiders’, ‘guarders’, ‘clutch tenders’, ‘nesters’, ‘external brooders’, and ‘internal livebearers’ (levels 1-9, 576 species). Principal component analysis (PCA) was performed on species with complete data for all five ecological metrics (382 species) using the function prcomp in the stats package in R.

### Construction of phylogenetic trees and other plots

Phylogenetic trees were constructed using ggtree^42^. Plots on diurnal/nocturnal transitions through time were generated with ggplot2 using data extracted from the ancestral reconstructions using custom code in R. The geological timescales were generated with code modified from the deeptime v1.0.1 package (https://zenodo.org/record/7709533). The function ggbiplot was used to generate PCA plots (https://github.com/vqv/ggbiplot). All other plots were generated using ggplot2 and final figure layouts were assembled in Affinity Designer (Serif Europe). All code used to perform analysis and generate figures is available online (https://github.com/maxshafer/fish_sleep).

## ACKNOWLEDGEMENTS

We thank members of the Shafer, Schier and Salzburger laboratories for discussion and advice, as well as F. Ronco for guidance regarding ancestral reconstructions. We thank A. Sawh and B. Chang for helpful comments on the manuscript. This work was supported by grants from the Swiss National Science Foundation (SNSF) to M.E.R.S. (SPARK 196313), A.F.S. (197837), and W.S. (208002), and from the Human Frontier Science Program (HFSP) to A.L.A.N (LT000400/2019-L). We also thank all of the previously published literature, and countless theses that enabled this study.

## AUTHOR CONTRIBUTIONS

M.E.R.S. and A.L.A.N. conceived and designed the study, and collected data from the literature. M.E.R.S. performed all analysis. M.E.R.S., A.L.A.N., A.F.S., and W.S. interpreted the results and wrote the manuscript. All authors read and approved the manuscript.

---

**Extended Data Figure 1.**
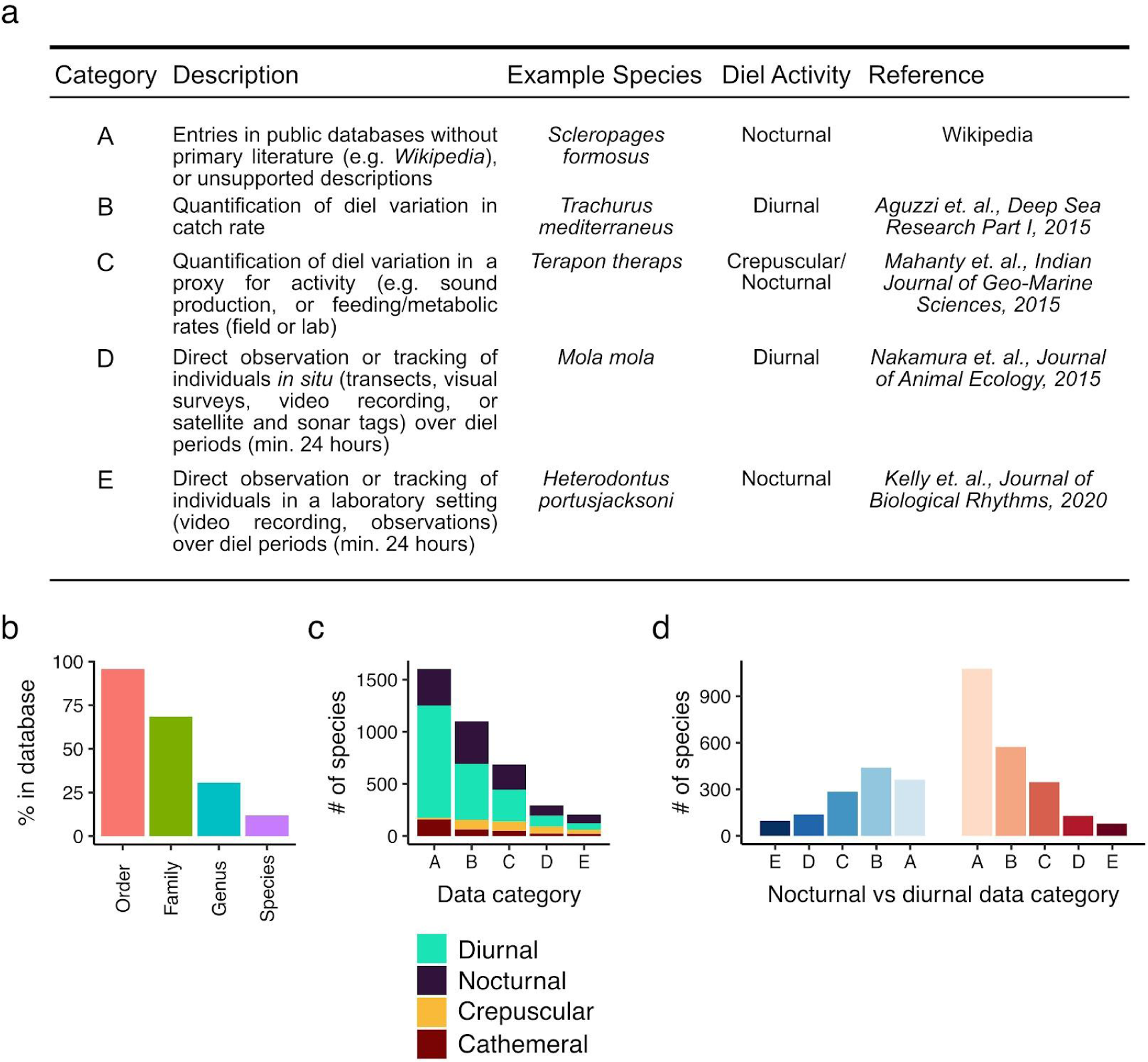
Literature survey and fish temporal activity pattern database statistics. **a**) Comparison of categories from public sources and primary literature corresponding to diel observations of fish species in our database, with example species with citations, along with their activity patterns. **b**) Coverage (as a percentage of taxonomies) within the database across taxonomic levels in Fishbase. **c**) Coverage (number of species) across categories and temporal niches. **d**) Coverage (# of species) across species coded as diurnal or nocturnal separated by category.^1^

**Extended Data Figure 2.**
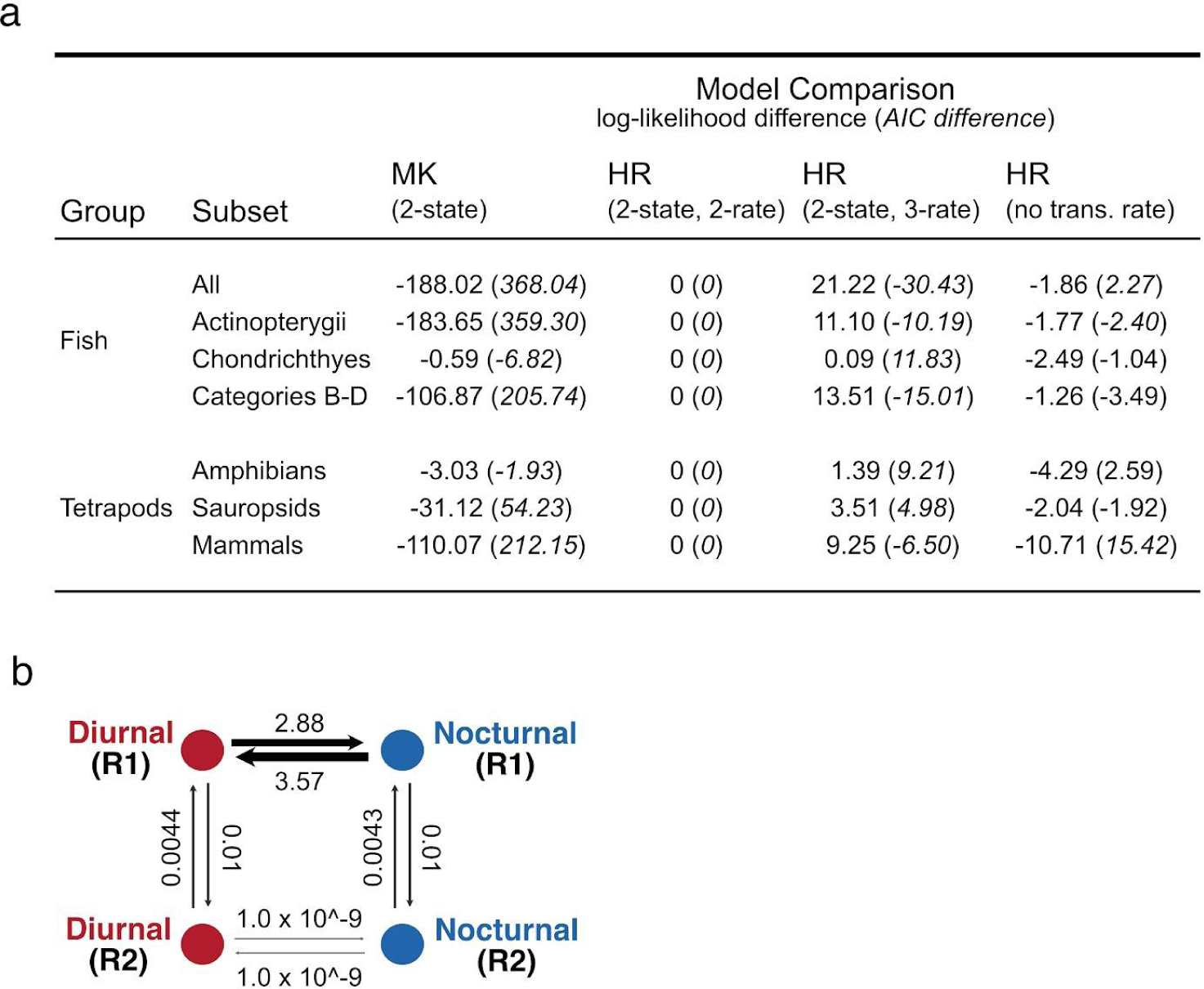
Comparison of models for the evolution of diel activity patterns. **a**) Comparison of model fits for different models of trait evolution using time-calibrated species trees with different topologies and subsets of species. Compared models include MK models with a single rate, models with 2 or 3 hidden rates (HR), or a model with a hidden rate with a transition probability of 0. Values are the log-likelihoods (or AIC values) minus the log-likelihoods (or AIC values) for the HR 2-state, 2-rate model. **b**) Plot showing the transition rates between states for the hidden rates model (HR) with two rates and two states for the dataset including both bony and cartilaginous fishes (“Fish - All”).

**Extended Data Figure 3.**
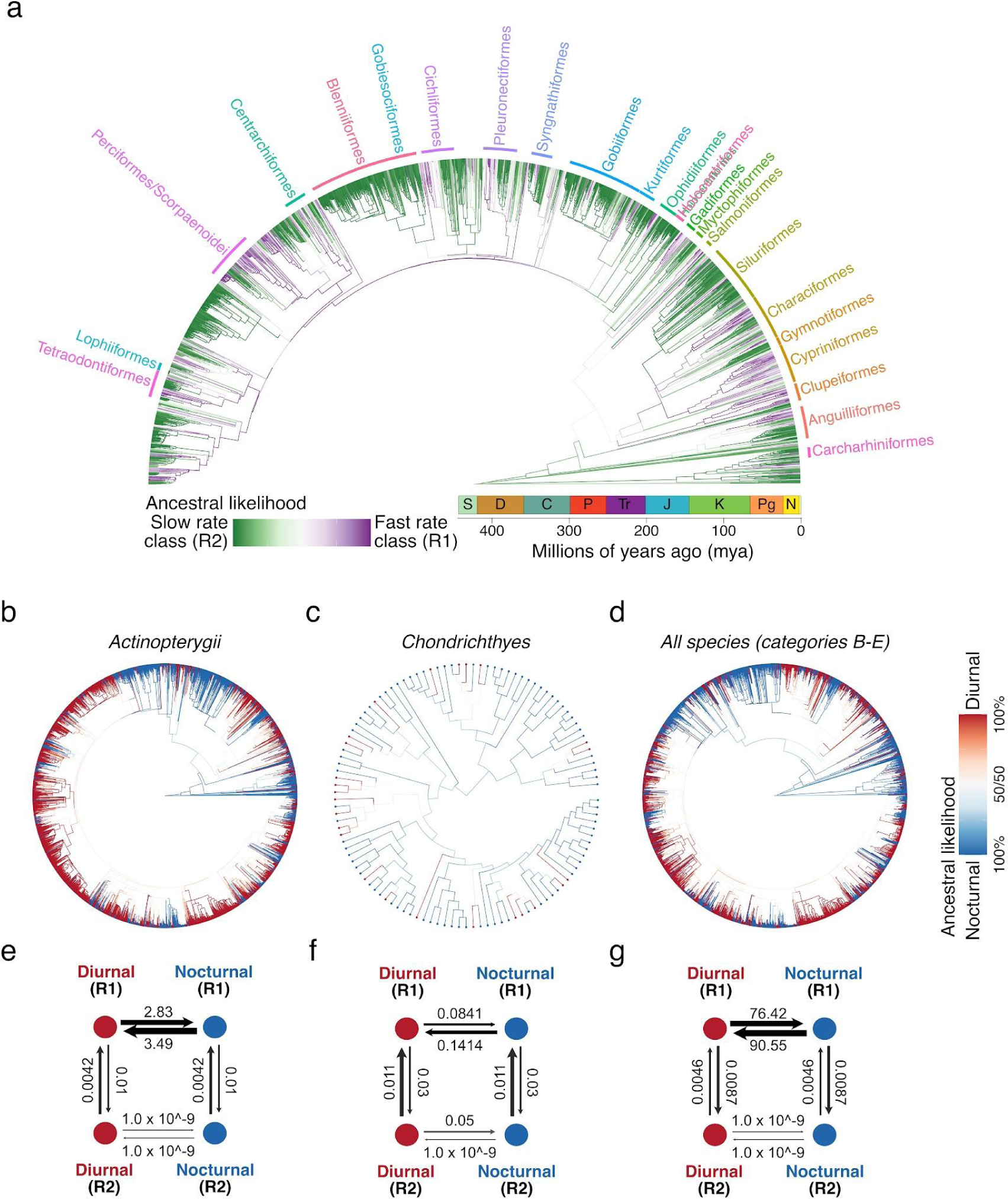
The evolution of temporal niche in fishes is best explained by the Hidden Rates Model. **a**) Marginal ancestral reconstruction of the slow and fast rate classes across all fish species. Marginal reconstructions and across only bony fish species (3401 species) (**b**) only cartilaginous fish species (113 species) (**c**), and all fish species from publications in categories B-E (2085 species) (**d**). Plots showing the transition rates between states for the hidden rates model (HR) with two rates and two states for bony fish species (3401 species) (**e**) only cartilaginous fish species (113 species) (**f**), and all fish species from publications in categories B-E (2085 species) (**g**).

**Extended Data Figure 4.**
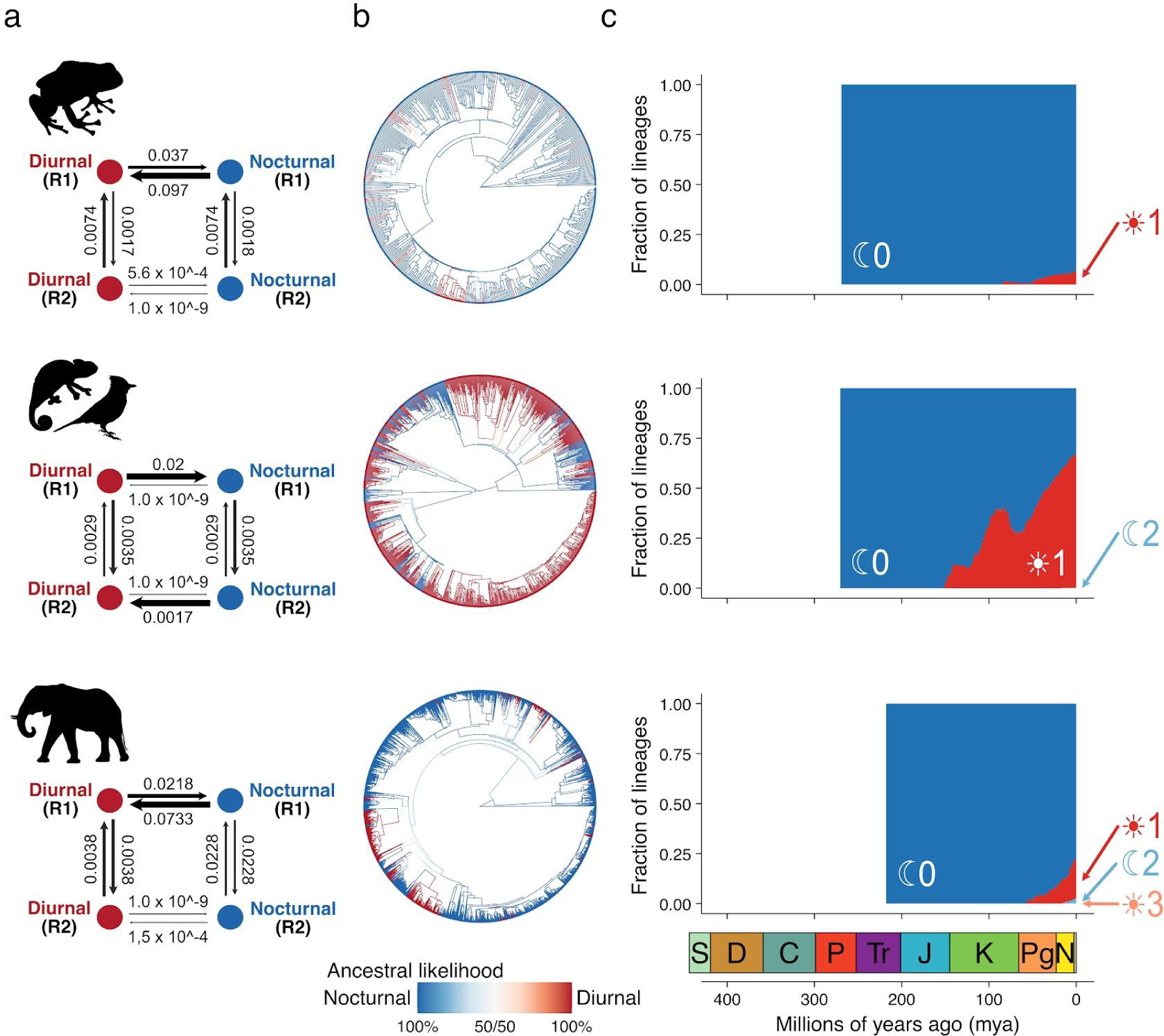
Ancestral reconstructions of diurnal/nocturnal behaviour across bony vertebrates. **a**) Rate plots for the best fitting models for the evolution of diurnal/nocturnal behaviour across amphibians, sauropsids and mammals. **b**) Marginal ancestral reconstructions of diurnal/nocturnal behaviour across amphibians (470 species), sauropsids (1096 species), and mammals (4544 species). **c**) The accumulation of lineages across time with a reconstructed history of 0, 1, 2, or 3 transitions between diurnal/nocturnal behaviour across amphibians, sauropsids, and mammals.

**Extended Data Figure 5.**
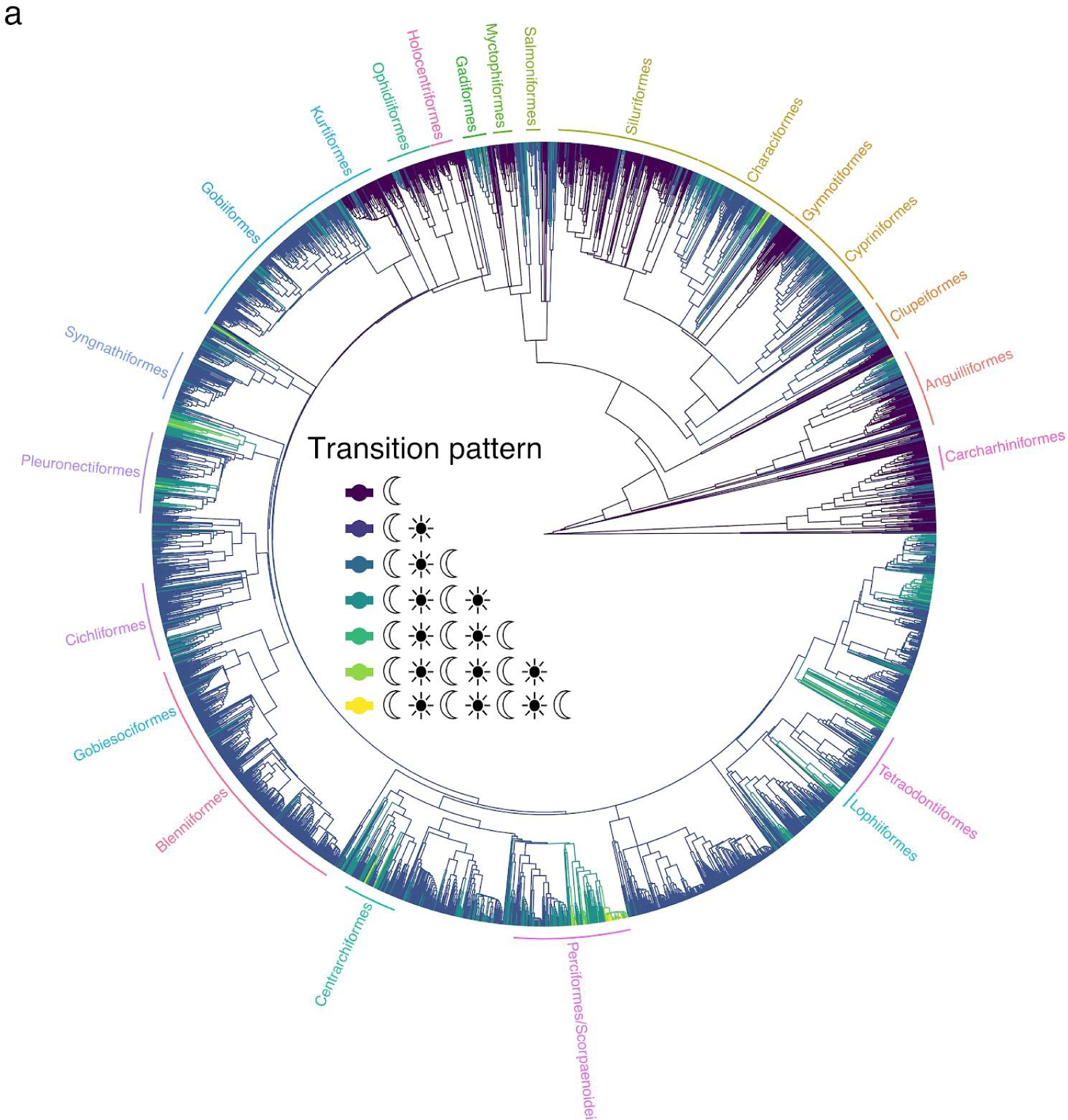
Ancestral reconstruction of the number of transitions in diurnal/nocturnal behaviour across bony fishes. **a**) Phylogenetic tree with nodes and branches coloured by the evolutionary history and number of transitions in diurnal/nocturnal behaviour. Phylogenetic orders are indicated by labels around the tree.

**Extended Data Figure 6.**
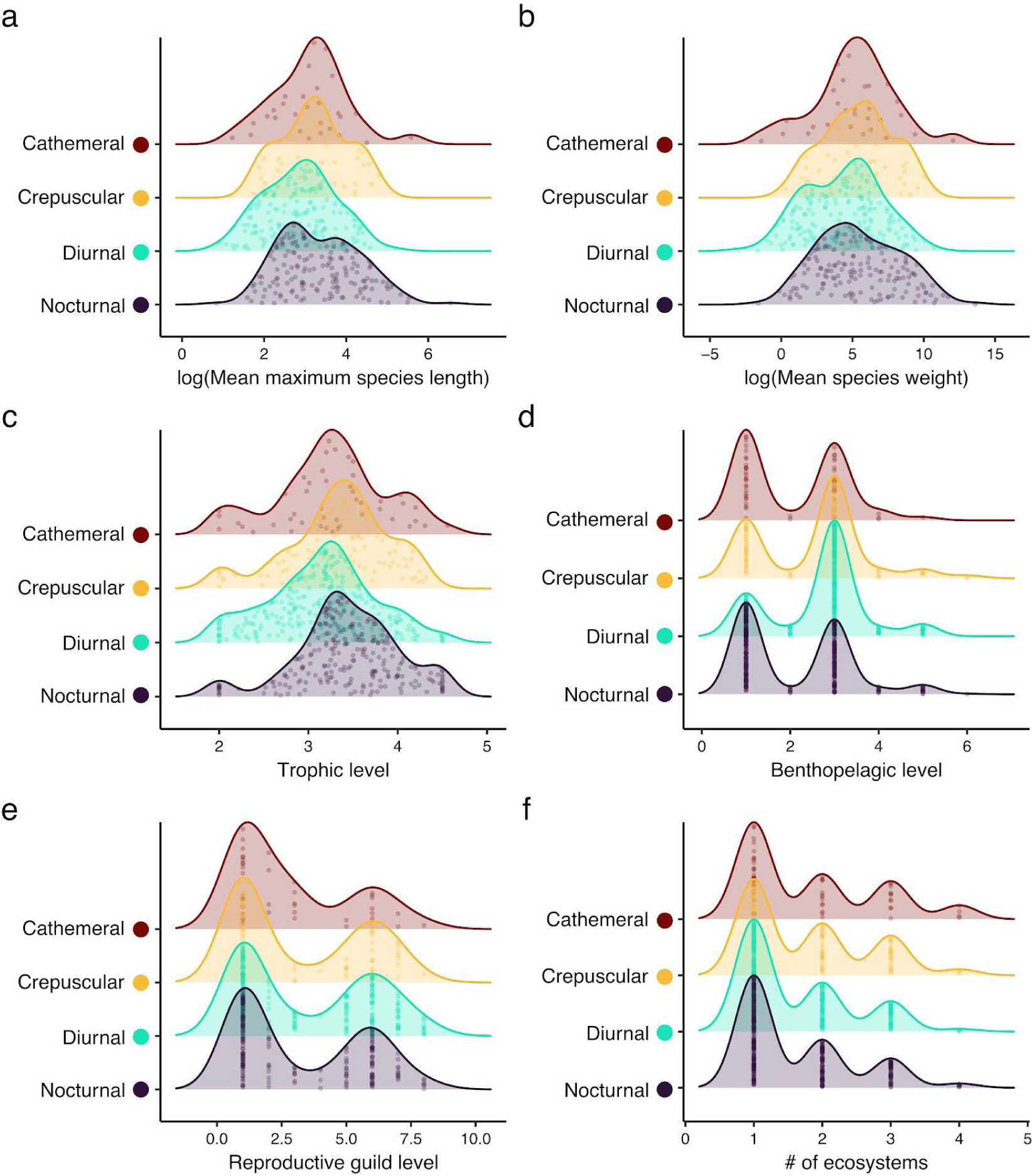
Distribution of ecological traits across nocturnal, diurnal, crepuscular, and arrhythmic fish species. Distribution across all fish species with recorded data on maximum length (**a**), mean weight (**b**), trophic level (**c**), benthopelagic axis (**d**), reproductive guild (**e**), and the number of occupied ecosystems (ecosystem breadth) (**f**).

**Extended Data Figure 7.**
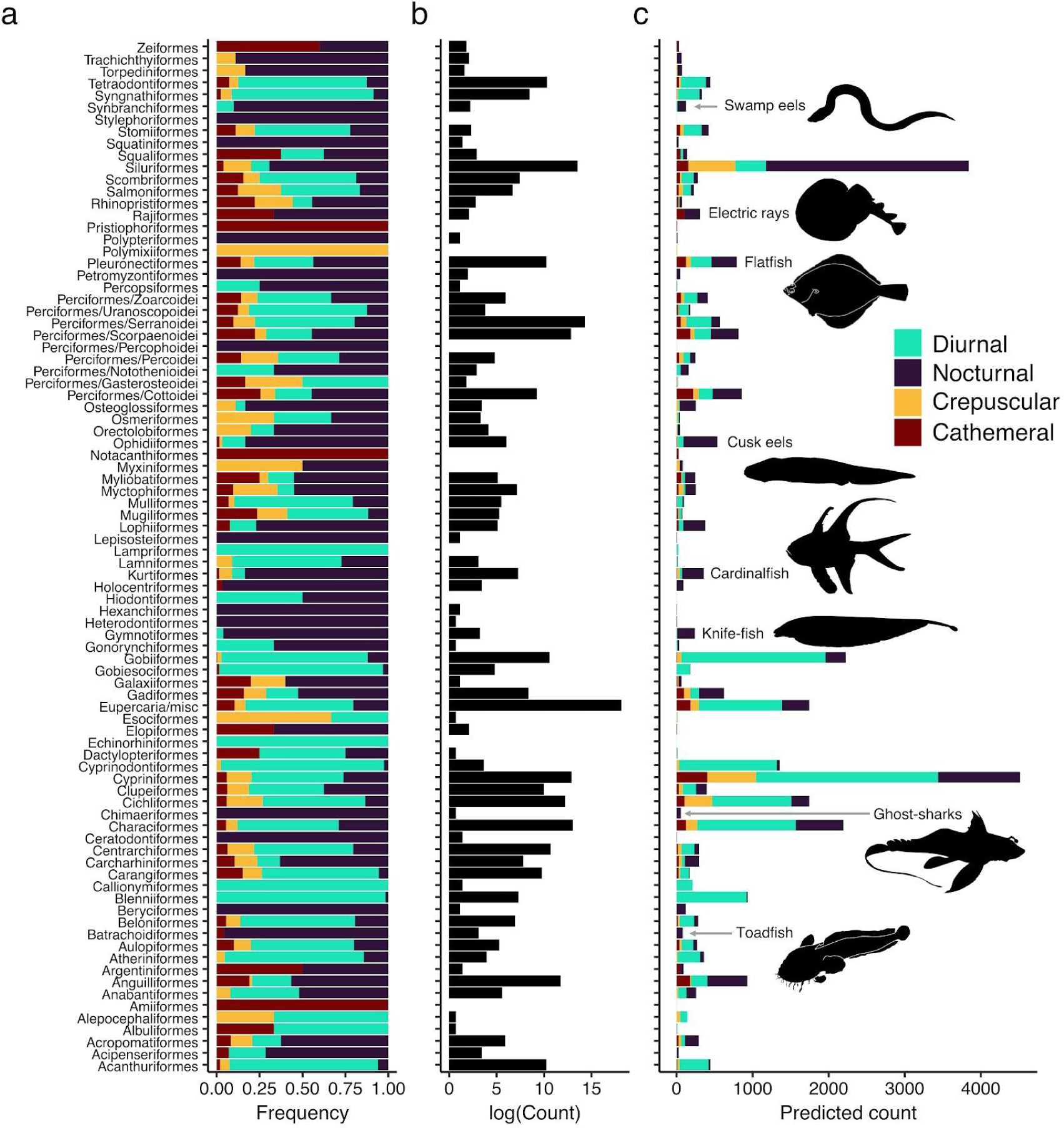
Predicted temporal niches across bony and cartilaginous fishes. **a**) Frequency of temporal niche for each taxonomic order of bony and cartilaginous fish. **b**) Total number of species for each taxonomic order of bony and cartilaginous fish. **c**) Predicted number of temporal niches for each taxonomic order of bony and cartilaginous fish.

**Extended Data Figure 8.**
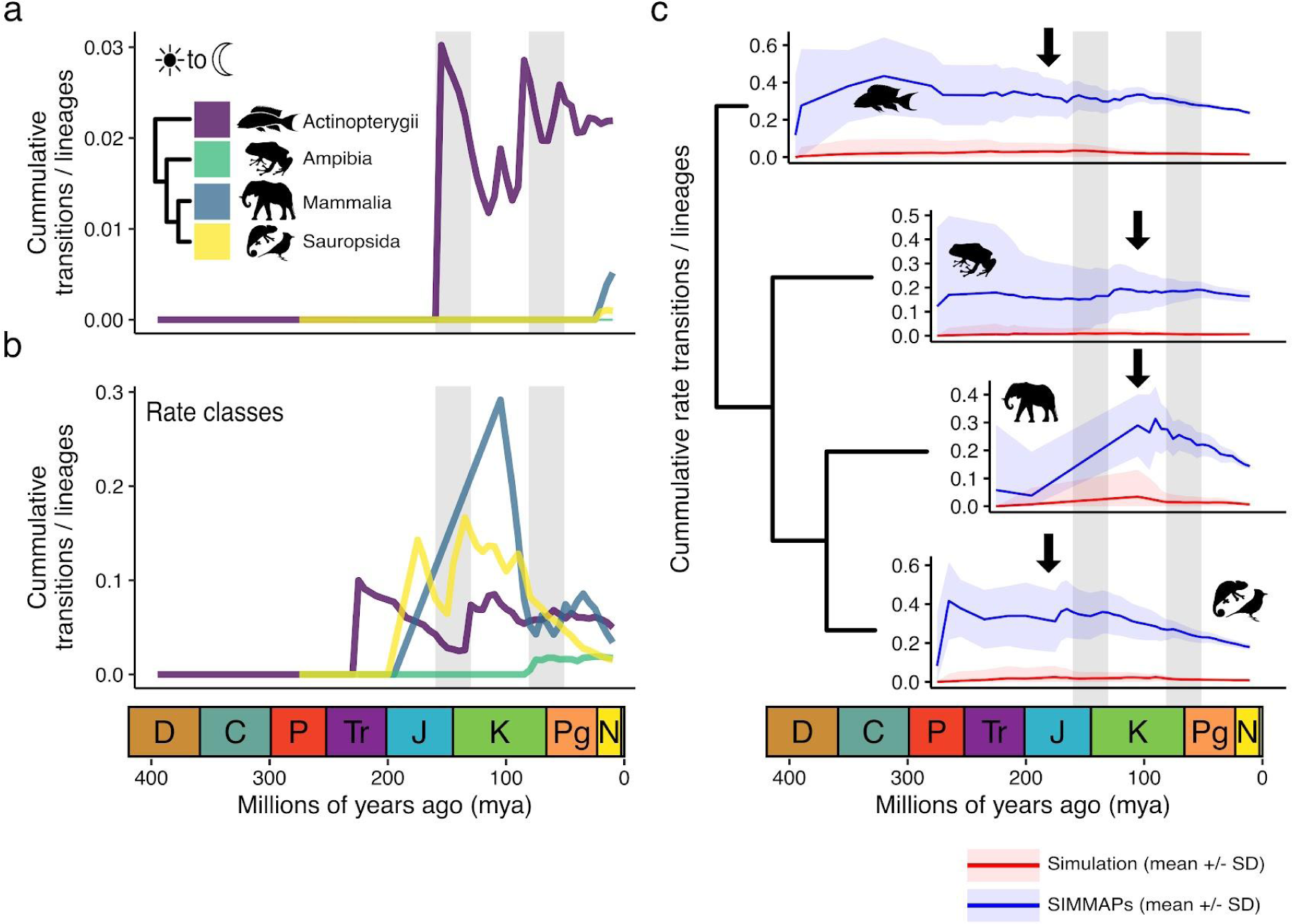
Transitions between rate classes suggest elevated rates prior to paleontological boundary periods. **a**) Cumulative number of transitions from diurnal to nocturnal activity patterns divided by the cumulative number of lineages per bony vertebrate group. Inset: the phylogenetic relationship between the groups. **b**) Cumulative number of transitions in rate classes (slow and fast) divided by the cumulative number of lineages per bony vertebrate group. **c**) Comparisons in transitions between rate classes from 500 simulations of the evolution of diurnal/nocturnal behaviour and 500 samples from the posterior distribution of our ancestral reconstructions (SIMMAPs) across bony vertebrate groups. Arrows indicate periods before geological boundaries when simulations and SIMMAPs differ. Dark lines are the means of simulations or SIMMAPs and shaded areas represent one standard deviation. Transitions are averaged across 5 million year bins.

**Extended Data Figure 9.**
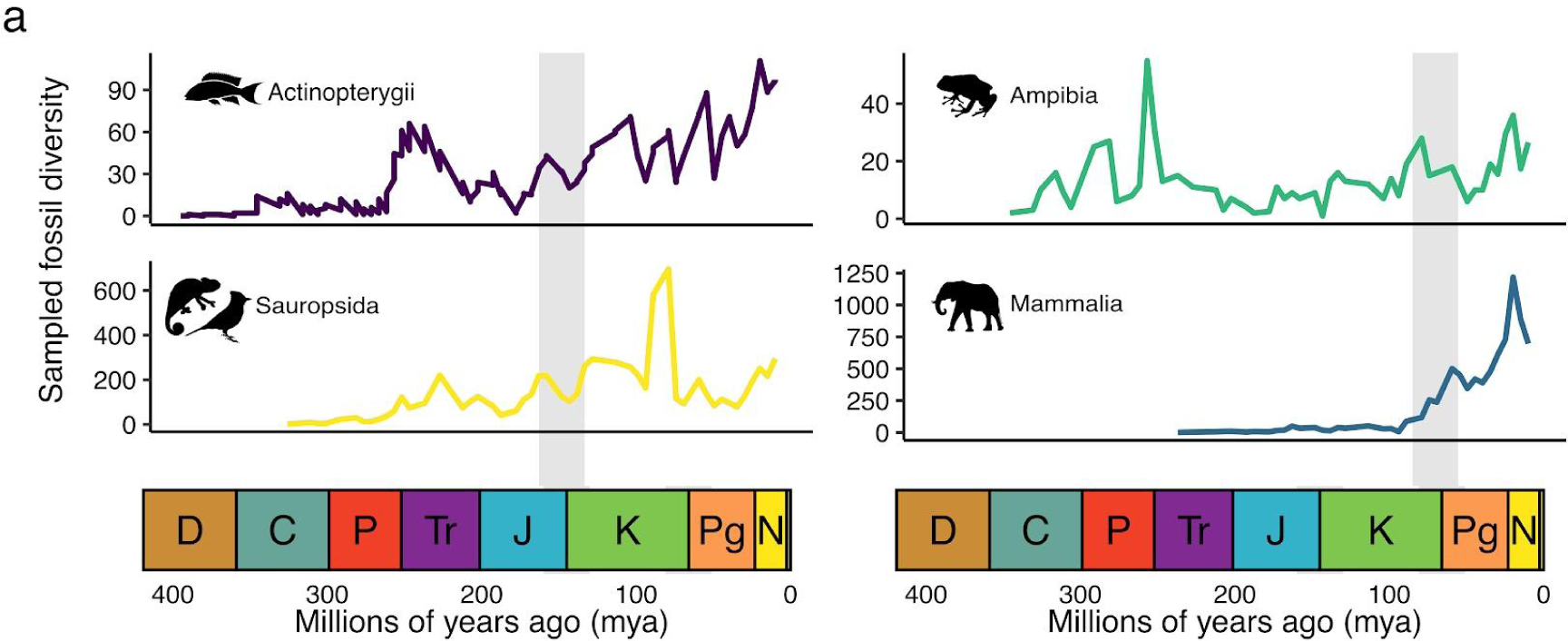
Fossil diversity of major bony vertebrate groups across geological time. **a**) Sampled fossil diversity for each of the major bony vertebrate groups before, during, and after the palaeontological boundary associated with each extinction bottleneck. Grey bars indicate bottleneck periods. Data on fossil records were extracted from the Paleobiology Database (https://paleobiodb.org/).

